# Brain connectivity dynamics: Multilayer network switching rate predicts brain performance

**DOI:** 10.1101/403105

**Authors:** Mangor Pedersen, Andrew Zalesky, Amir Omidvarnia, Graeme D. Jackson

**Affiliations:** The Florey Institute of Neuroscience and Mental Health, The University of Melbourne, VIC, Australia; Department of Psychiatry, Melbourne Neuropsychiatry Centre, The University of Melbourne, VIC, Australia; Melbourne School of Engineering, The University of Melbourne, VIC, Australia; Department of Neurology, Austin Health, Melbourne, VIC, Australia

**Keywords:** Network switching, Dynamic connectivity, fMRI, Brain and behaviour

## Abstract

Large-scale brain dynamics measures repeating spatiotemporal connectivity patterns that reflect a range of putative different brain states that underlie the dynamic repertoire of brain functions. The role of transition between brain networks is poorly understood and whether switching between these states is important for behavior has been little studied. Our aim here is to model switching between functional brain networks using multilayer network methods and test for associations between model parameters and behavioral measures. We calculated time-resolved functional MRI (fMRI) connectivity from one-hour long data recordings in 1003 healthy human adults from the Human Connectome Project. The time-resolved fMRI connectivity data was used to generate a spatiotemporal multilayer modularity model enabling us to quantify *network switching* which we define as the rate at which each brain region transits between different fMRI networks. We found i) an inverse relationship between network switching and connectivity dynamics –defined as the difference in variance between time-resolved fMRI connectivity signals and phase randomized surrogates–; ii) brain connectivity was lower during intervals of network switching; iii) brain areas with frequent network switching had greater temporal complexity; iv) brain areas with high network switching were located in association cortices; and v) using cross-validated Elastic Net regression, network switching predicted inter-subject variation in working memory performance, planning/reasoning and amount of sleep. Our findings shed new light on the importance of brain dynamics predicting task performance and amount of sleep. The ability to switch between network configurations thus appears to be a fundamental feature of optimal brain function.

## Introduction

Functional MRI (fMRI) has significantly enhanced our knowledge about human brain function (Bandettini, Wong, Hinks, Tikofsky, & Hyde, 1992; Kwong et al., 1992; Ogawa et al., 1992), especially in recent years when fMRI data has been used to quantify the brain as a complex network (Rubinov & Sporns, 2010; Bullmore & Sporns, 2009). Although fMRI-based network analyses have led to several new insights into the spatial and temporal nature of large-scale brain network activity (Hutchison et al., 2013; Preti, Bolton, & Van De Ville, 2017), many early fMRI network studies treat spatial and temporal information as separate entities, meaning that brain regions are not interconnected across time and space.

Multilayer network analysis (Muldoon & Bassett, 2016; De Domenico, 2017) is a novel graph-theoretic model of networks where nodes are connected across time and space. Multilayer networks can be decomposed into modules that span time and space using a multilayer modularity algorithm (Mucha, Richardson, Macon, Porter, & Onnela, 2010) that estimates the spatiotemporal segregation of nodes forming a subset of non-overlapping modules or networks. This multilayer modularity model has a major advantage over other time-resolved fMRI connectivity methods as it provides a ‘temporal link’ or connectivity between adjacent time points. In other words, the multilayer modularity model allows us track and quantify temporal changes of each node, and also, when they ‘switch’ between different module or network assignments (Bassett et al., 2013). Network switching is defined as *the rate at which a brain region transitions between different functional networks.* Note this measure has previously been called node flexibility as proposed by Bassett and others (Bassett et al., 2013, 2011), however, we prefer the term node switching which has also been recently used by Bassett’s group (Telesford et al., 2017). Despite multilayer modularity being a relatively new technique, a series of studies suggest that network switching is associated with learning of simple motor tasks (Bassett et al., 2011), attention (Shine, Koyejo, & Poldrack, 2016), executive function (Braun et al., 2015), fatigue (Betzel, Satterthwaite, Gold, & Bassett, 2017) and depression (Zheng et al., 2017).

These studies suggest that multilayer modularity has an underlying neurobiological basis; however, it remains unknown whether network switching is correlated with the dynamics, or variance, of fMRI connectivity time-series, and whether network switching occurs during time-periods of high or low network connectivity and complexity. It is important to enhance our understanding about switching and dynamics of fMRI connectivity in order to reconcile how state changes and switching of networks (topology analysis) may relate to statistical dynamics theory (signal analysis). In an attempt to address these non-trivial questions and gaps in the literature, we investigate network switching in a multilayer modularity model using time-resolved fMRI connectivity data from 1003 healthy adults provided by the Human Connectome Project (Van Essen et al., 2013). We hypothesize that fMRI-based network switching, and connectivity dynamics are intrinsically correlated. Given that network switching is likely to be a ‘strenuous’ event for the brain, we also hypothesize that network switching is associated with changes in fMRI complexity and connectivity. Lastly, we hypothesize that network switching is correlated with cognitively demanding behavioral tasks.

## Results

Time-resolved fMRI connectivity was estimated with correlation-based sliding window analysis from 25 brain nodes (*N* – all brain nodes were derived from an Independent Components Analysis across all subjects; all nodes are visualized in Supplementary Figure 1), and 4800 time-points (≈ 1 hour of data – concatenating 4×14.4 min sessions of fMRI data). We used a window length of 100 seconds (139 timepoints), and a window overlap of 0.72 seconds (1 time-point). This resulted in 4661 ‘time-windows’ (W) forming three-dimensional correlation coefficient matrices for each subject with a size of N×N×W. This time-resolved fMRI connectivity data was used as an input to the multilayer modularity model, which was an iterative Louvain community detection algorithm with uniform ordinal temporal coupling between adjacent time-points (Mucha et al., 2010). The temporal coupling strength of this model is governed by an *ω* parameter, whereas the topological resolution of modules is governed by a *γ* parameter. Low/high *ω* provides weak/strong temporal coupling between adjacent time-points, whereas low/high *γ* give few/many spatial modules. The most commonly used parameters in this multilayer modularity algorithm are *ω* = *γ* = 1. However, to ensure that our results are not affected by a specific spatial and temporal parameter, we calculated multilayer modularity across a range of parameter values including *γ* = [0.9, 1, 1.1]; *ω* = [0.5, 0.75, 1], previously found to have strong spatiotemporal modularity (11). The output of the multilayer modularity model was a two-dimensional array (N W) containing integer values outlining spatiotemporal nodal network assignments. We then calculated nodal network switching as the proportion of layers (time windows) in which a node’s network assignment changes (see Figure 1, for an overview).

**Figure 1.**
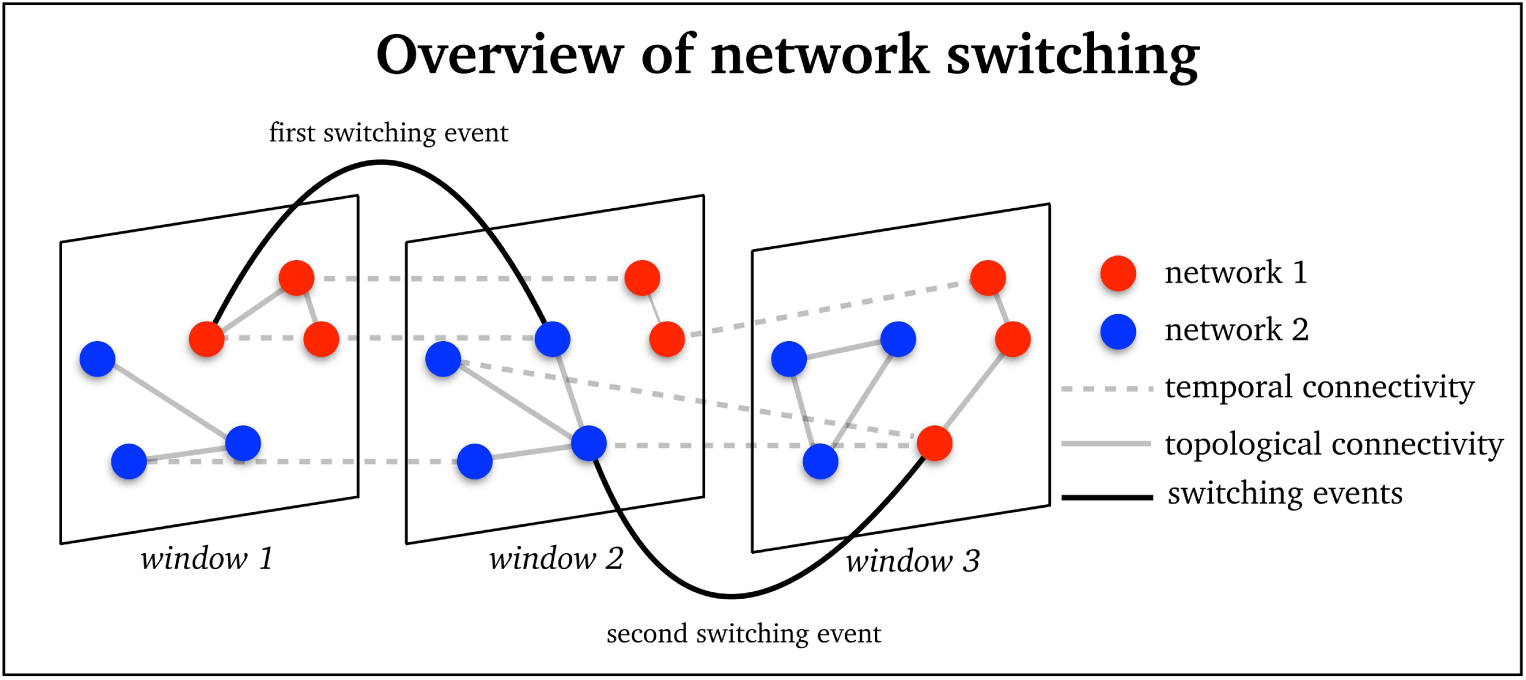
An overview of network switching within a multilayer modularity network with 6 nodes and 3 time-windows (window 1, window 2 and window 3) and 2 modularity partitions (red = network 1; blue = network 2). This example shows 2 switching events exemplified when node changes between red and blue colors between time-points (solid black line between time-points). Solid grey lines correspond to within-layer, or topological, connectivity. Dashed grey lines correspond to between-layer, or temporal, connectivity.

### Switching and dynamics are inversely related

Firstly, we assessed whether network switching was related to dynamic connectivity. Dynamic connectivity was defined as the difference between the standard deviation of all fMRI sliding-windows connection-pairs and the standard deviation of 500 phase randomized surrogates (corresponding to the null hypothesis of an absence of any dynamics) (Prichard & Theiler, 1994). An uncorrected p-value was assigned to the standard deviation value of each fMRI sliding-window connection pair corresponding to its relative ‘rank’ compared to the 500 randomized surrogates. For example, an fMRI connection-pair will have an uncorrected p-value of 0.002 if it has a greater standard deviation value than 499 of the 500 randomized surrogates (calculated as 1-*rank*/total number of variables, where *rank* = 500 out of 501 total variables). Statistical significance of connectivity dynamics was then determined using a false discovery rate (Benjamini & Hochberg, 1995) correction with threshold of *q* = 0.05, over all uncorrected p-values. In order to reduce dynamic connectivity information from the level of connection-pairs ([*N*(*N* −1)]*/*2) = 300) to nodes (*N* = 25), we calculated the binary sum of all significant dynamic connections associated with each node. This resulted in a nodal degree metric quantifying the number of dynamic connections associated with each node.

We found that network switching was inversely correlated with fMRI-based connectivity dynamics. Averaged over all subjects, the Spearman’s correlation between nodal network switching and dynamics ranged between *ρ* = −0.49 and - 0.52, across all *ω* and *γ* parameters (Figure 2A – *ω* and y = 1 shown). Averaged over all brain nodes, the Spearman’s correlation between subject-level network switching and dynamics ranged between *ρ* = −0.51 and −0.55, across all *ω* and *γ* parameters (Figure 2B – *ω* and *γ* = 1 shown).

**Figure 2.**
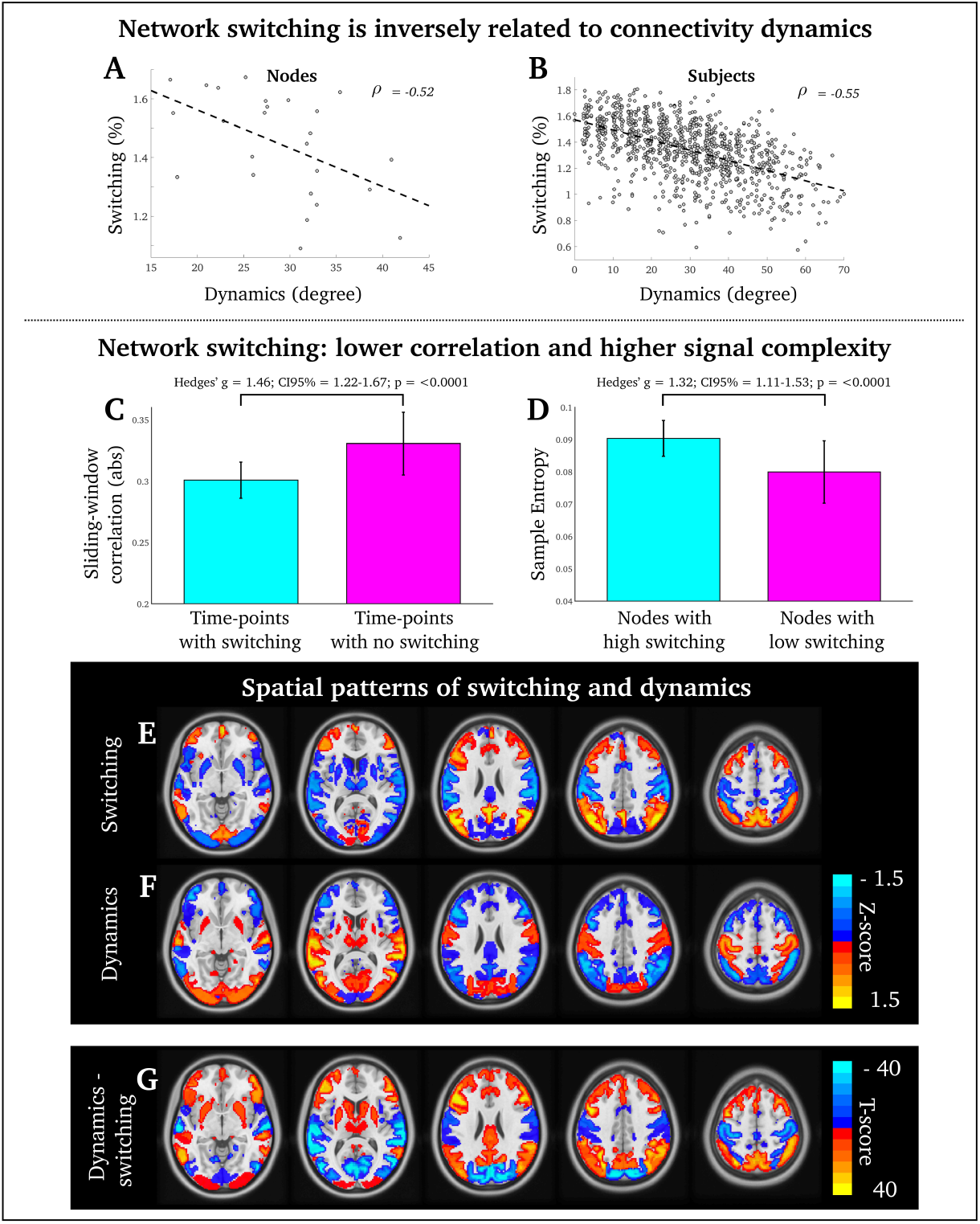
Scatter-plot between network switching and connectivity dynamics A) Each data-point denotes a single node, aver-aged across all subjects; B) Each data-point denotes a single subject, averaged across all nodes; C) During time-windows with network switching (cyan color), nodes display significantly lower absolute sliding-window correlations than time-windows with no switching (magenta color). D) Average Sample Entropy in the 5 nodes with most network switching (cyan color) was significantly higher than the 5 brain nodes with lowest network switching (magenta color). E) Network switching was high in association cortices and low in primary cortices. F) Dynamic connectivity was high in primary cortices and low in association cortices. G) Paired t-test difference between the 25 nodes in E and F. Data for all 25 brain nodes were normalized into z-scores so ensure both connectivity dynamics and switching values were scaled equally and appropriate for univariate t-test analysis.

In line with our prior hypothesis, topological switching and signal dynamics of networks are (inversely) correlated. This finding raises the possibility that the switching brain is associated with an overall reduction of brain connectivity, here sliding-window correlations. To further test this assumption, we calculated the average correlation coefficient of each sliding window correlation matrix corresponding to time-widows when brain regions switch between networks versus time-windows when brain regions do not switch between networks. Given that variation in *ω* and *γ* had a negligible impact, we henceforth only consider *ω* and *γ* = 1.

### Switching is frequent when network connectivity is low

The (absolute) average sliding-window correlation coefficient of all possible pair-wise correlations between nodes was significantly lower during time-windows when nodes switch between networks (i.e., between two neighboring layers when the ‘switch’ occurred), compared to time-windows when nodes do not switch between networks (Hedges’ *g* effect size = 1.46; 95^*th*^ percentile confidence interval = 1.22-1.67; df = 2004; p <0.0001 – Figure 2C). This suggests that network switching occurs during periods of low network connectivity. Given that this analysis was conducted in the topological domain, we next wanted to elucidate whether temporal complexity was also affected by network switching. To this end, we calculated entropy (signal complexity) of brain nodes that switch networks most frequently versus brain nodes that switch networks least frequently.

### Association between signal complexity and switching

We found significantly higher sample entropy (Richman & Moorman, 2000) (parameter values were *M* = 2; *r* = 0.2 times the standard deviation of signals) values of sliding-window correlation time-series in the five brain regions with highest rate of network switching (superior parietal lobule, precuneus, left frontoparietal lobe and right frontoparietal lobe and right intraparietal sulcus), compared to the five brain regions with lowest rate of network switching (secondary visual cortex, superior temporal lobe, primary motor cortex and left cerebellum and right cerebellum). Hedges’ *g* effect size = 1.32; 95^*th*^percentile confidence interval = 1.11-1.53; df = 2004; p <0.0001 (Figure 2D). This finding suggests that network switching is associated with temporally complex fMRI connectivity signals.

Insofar, our results suggest that i) switching and dynamics are negatively correlated, ii) switching time-windows have lower correlations in the topological domain and iii) frequently switching nodes have greater complexity in the temporal domain. Following this, we next sought to localize which cortical and subcortical regions switch the most.

### Switching is most frequent in association cortex

We observed a divergent spatial pattern between network switching and functional dynamics. Higher switching was observed in association cortex compared to primary cortex (hot colors in Figure 2E) whereas the converse pattern was evident for connectivity dynamics (hot colors in Figure 2F). In Figure 2G, we report results from a paired t-test outlining nodal differences between switching and dynamics. In total, 24 of 25 brain nodes were statistically different between switching and connectivity dynamics after correcting for multiple comparisons (False Discovery Rate, *q* = 0.05). The only brain region not statistically significant between switching and dynamics was the secondary visual cortex (see node 4 in Supplementary Figure 1).

In light of these spatial network findings (Figure 2 E-F), we next aimed to confirm that the multilayer modularity algorithm delineated spatial modules conforming to well-known resting-state networks. A module consensus map was generated using a group-averaged agreement matrix that contains probability values [0,1] denoting the number of times node-pairs share the same module divided by the number of possible times two nodes can share the same module. By computing a Louvain clustering algorithm of the group-averaged agreement matrix (Lancichinetti & Fortunato, 2012), we obtained five well-known modules, or networks, common to all subjects including: i) primary sensory cortices, ii) secondary sensory cortices, iii) salience/subcortical network, iv) fronto-parietal network and v) default mode network (see Supplementary Figure 2).

### Relationship between switching and behavior

Lastly, we aimed to test whether switching predicted inter-individual variation in behavior and task performance. Given that the Human Connectome Project offers a wealth of behavioral information (Barch et al., 2013), we wanted to use a data-driven regression approach without any prior bias. To this end, we included 50 behavioral variables comprising behavioral domains such as cognition (working memory, attention, executive functioning, planning, reasoning and gambling), social functioning, personality traits, physical function and sleep. Although we used a data driven regression approach, we hypothesized that the subset of tasks important for ‘higher-order’ or ‘frontal lobe’ function would be most important for efficient brain network switching. These cognitive tasks include: i) the Flanker task (cognitive inhibition); i) card sorting task (cognitive flexibility); iii) processing speed (general cognitive ability); iv) N-back task (working memory); and v) relational task (planning and reasoning).

We used Elastic net regression (Zou & Hastie, 2005) to test whether any of the 50 behavioral domains (independent variables) predicted whole-brain averaged network switching (dependent variable). Elastic net regression is well suited to data-driven regression analysis as it provides a sparse output by removing all behavioral data deemed to be unrelated to network switching. Elastic net is governed a regularization parameter *λ* that alters the sparsity and variability of the regression model. The regularization parameter was determined with 10-fold cross validation (Lachenbruch & Mickey, 1968). The minimum mean square error (0.028) was achieved with a regularization parameter *λ* = 0.023 (see Supplementary Figure 3). At this value, behavioral data accounted for ≈ 3% of the total variance of fMRI network switching data (r^2^ = 0.029, defined as 1-[residual sum of squares/total sum of squares] of the regression model). This r^2^ value was significantly higher than expected due to chance (p <0.001, compared to r^2^ estimates from 500 randomly generated Elastic Net regressions).

**Figure 3.**
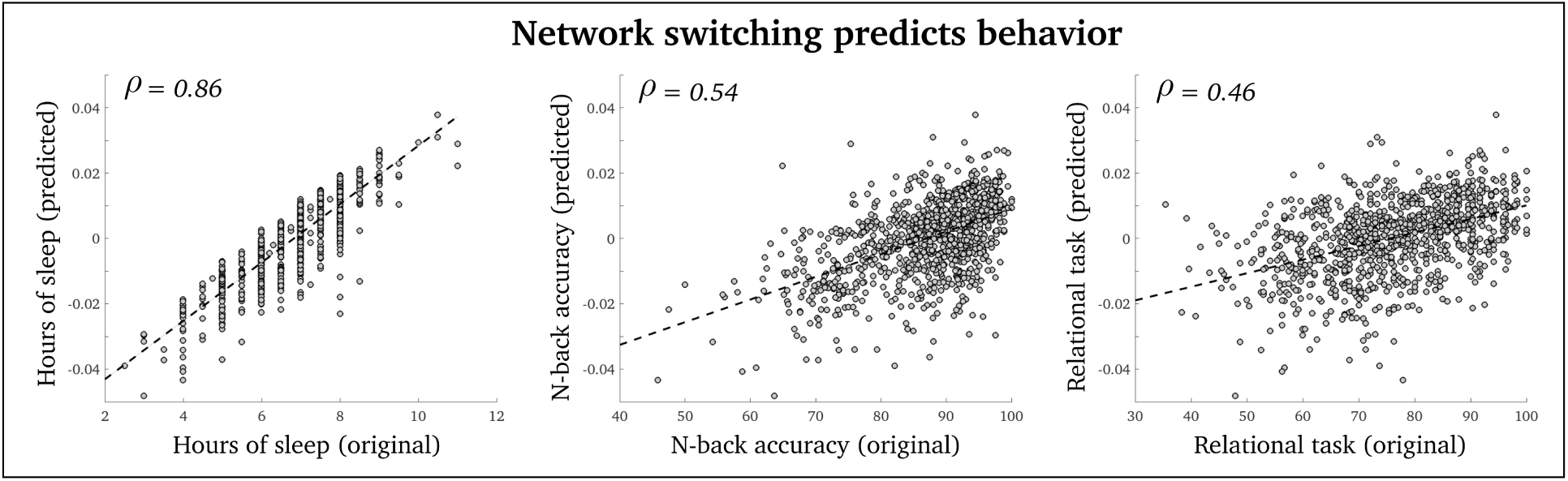
Scatterplots between Elastic net predicted and rescaled behavioral variables (y-axes) and original values (x-axes). Stippled lines are the best linear fit between predicted and original values. Each data-point denote a subject.

The Elastic net regression result at *λ* of 0.023 showed that 3 of 50 behavioral domains were weakly, but significantly, related to network switching (note that Elastic net regression *α* values were zero for all other 47 measures): i) number of hours of sleep the night before the MRI scan (Spearman’s correlation between switching and hours of sleep was *ρ* = 0.14, p <0.0001; Elastic net regression *β* was 0.071; the Spearman’s correlation between original and predicted data was *ρ* = 0.86 – Figure 3 left); As hypothesized, higher-order functions were also correlated with switching: ii) accuracy of N-back task using the average accuracy score from the 0-and 2-back tasks, important for working memory (Spearman’s correlation between switching and N-back task was *ρ* = 0.11, p <0.0001; Elastic net regression *α* was 0.036; the Spearman’s correlation between original and predicted data was *ρ* = 0.54 – Figure 3 middle); and iii) relational task involved in planning and reasoning (Spearman’s correlation between switching and relational task was *ρ* = 0.11, p <0.0001; Elastic net regression *α* was 0.017; the Spearman’s correlation between original and predicted data was *ρ* = 0.46 – Figure 3 right).

## Discussion

By leveraging the information-rich brain imaging dataset (approximately one-hour fMRI recordings from 1003 subjects) provided by the Human Connectome Project (Van Essen et al., 2013) we found that fMRI network switching was inversely correlated with the dynamics of fMRI connectivity (Figure 2 A-B), with most prominent network switching in association cortices (Figure 2 E). Although it is unlikely that these high switching nodes (e.g., bilateral frontal and parietal cortex) do not exhibit dynamic activity, they do not conform to our current metric of standard deviation as a proxy of dynamic connectivity, suggesting they have unchanging statistical properties over time. We also found that brain nodes switch between networks during time-windows with low network connectivity (Figure 2C), and high-switching nodes were more ‘temporally complex’ –estimated with Sample Entropy– compared to low-switching nodes (Figure 2D). Switching is known to increase in systems with high entropy or information load (Amigó, Kloeden, & Giménez, 2013). We consequently hypothesize that our observed relationship between brain network switching and high entropy/low network connectivity may be related to increased information load imposed on specific brain regions, especially those located within the association cortex (e.g., bilateral fontal and parietal cortex) known to integrate information between a range of different networks (Van Den Heuvel & Sporns, 2011).

Network switching also predicted inter-subject behavior, using data-driven Elastic net regression (Figure 3). Even though we included a range of behavioral variables such as personality traits, physical performance, emotion and well-being, behaviors that tap into association cortex function were predicted by network switching, including working memory and planning/reasoning (Courtney, Petit, Maisog, Ungerleider, & Haxby, 1998). We noted a resemblance between our spatial network switching map displayed in Figure 2D and previous task-based fMRI studies of working memory tasks (Wager & Smith, 2003), with prominent bilateral frontal and parietal cortices important for higher-order cognitive functions. Notably, network switching predicted the amount of sleep that participants had the night before the MRI scan. As all participants were instructed to keep their eyes open during the whole scan, falling asleep in the scanner cannot explain this finding. Sleep impacts on the same domains of brain performance as seen in this study (Alhola & Polo-Kantola, 2007) and findings by (Betzel et al., 2017) also suggest that fatigue may be an important ‘driver’ for network switching. Taken together, this suggests that the impact of sleep deprivation on cognitive performance may be mediated through its e?ect on brain network switching.

The computational complexity of multilayer network analyses was a limitation of our study as high spatial and/or temporal resolution of multilayer networks results in very large networks as they are connected in both time and space ([(N×W)(N×W)] ≈ 10^10^ data points per subjects in this study). Given that we needed to include many time-windows (here, 4800 time-windows) to statistically detect fMRI dynamic connectivity –this was previously demonstrated by (Hindriks et al., 2016), and further validated in this study as seen in Supplementary Figure 4–, we needed to keep the spatial resolution of fMRI data rather coarse (here, 25 brain nodes). We hope that more efficient computational methods will be developed in the future to enable assessment of multilayer network modularity in fMRI data with high spatial and temporal resolution.

It is worth noting that the main limitation of only having 25 network nodes is the so-called ‘resolution limit’ of modularity algorithms. This means that at some point there are insufficient number of nodes that can converge into segregated and non-overlapping modules. The resolution limit did not appear to be a problem in the current study as nodes were readily sub-divided into five well-known ‘resting-state networks’ across subjects associated with relatively high modularity scores (Q = ≈ 0.6) serving as a quality function of the obtained multilayer modularity partitions (Supplementary Figure 2).

## Methods and Materials

### Subjects, fMRI data and processing

We used resting state fMRI data from 1003 healthy adults from the Human Connectome Project (Van Essen et al., 2012) (female subjects = 534/1003; male subjects = 469/1003) and Institutional Review Board approval was considered unnecessary for the current study. fMRI recon r177+r227 data was used and subject were between ages of 22 and 35 years. fMRI parameters included: echo time = 33.1 ms; field of view = 208×180 mm^2^; number of slices = 72; voxel size = 2 mm^3^ and flip angle = 52 ^°^. We used four fMRI scans for each subject (14.4 minutes per scans where subjects were instructed to keep their eyes open). At a repetition time of 0.72 seconds there were 1200 time-points in each scan - we concatenated all four scans into continuous fMRI time-series comprising 4800 time-points.

The fMRI data of each subject was preprocessed by the Human Connectome team with echo planar imaging gradient distortion correction, motion correction, field bias correction, spatial transformation and normalization into a common Montreal Neurological Institute space (Glasser et al., 2013), and artefact removal using Independent Component Analysis FIX (Salimi-Khorshidi et al., 2014). A group-level Independent Component Analysis was used to define the 25 brain nodes of interest, common across all subjects. We additionally filtered the fMRI data between frequencies of 0.01 and 0.1 *Hz*.

### fMRI correlation-based sliding-windows analysis

We used a Pearson’s correlation-based sliding window analysis to estimate time-resolved fMRI connectivity. Pearson’s correlation coefficient between two fMRI time-series (Zalesky, Fornito, Cocchi, Gollo, & Breakspear, 2014; Keilholz, Magnuson, Pan, Willis, & Thompson, 2013; Pedersen, Omidvarnia, Zalesky, & Jackson, 2018) *X*[*t*] and *Y*[*t*] of length *T* is written as:

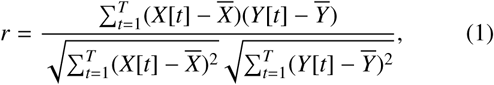

where 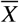 and 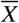 denote sample means and *r* ranges from −1 (full anti-correlation) to 1 (full correlation). Here, the pair-wise correlation coefficient between 25 brain regions-of-interest were calculated based on a fixed window length consisting of 139 fMRI time-point (100 seconds), which satisfies the 1/*f*_0_ wavelength criterion for a minimum cut-off frequency of 0.01*Hz* (Leonardi & Van De Ville, 2015; Zalesky & Breakspear, 2015). Windows were shifted with single-frame increments resulting in a total number of 4661 windows. We tapered each correlation-based window with a Hamming function to mitigate edge-artefacts of the windows and attenuate potentially noisy signals.

### Multilayer modularity and network switching

To quantify spatiotemporal network switching we used an iterative and ordinal Louvain algorithm to track network function over time (Mucha et al., 2010) (implemented with codes from Lucas G. S. Jeub, Marya Bazzi, Inderjit S. Jutla, and Peter J. Mucha, ‘A generalized Louvain method for community detection implemented in MATLAB,’ http://netwiki.amath.unc.edu/GenLouvain (2011-2016)). Modularity is quantified by *Q* ranging from 0 (low network segregation) to 1 (high network segregation). This measure is governed by *γ* and *ω* parameters, which determine the strength of topological and temporal connectivity, respectively. Multilayer modularity is written as following:

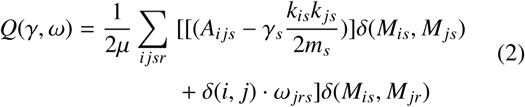

*A*_*ijs*_ is the sliding window correlation matrix between node *i* and *j*, for time-point *s* whereas 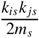 (*k* = node degree at time-point *s*; *m* = sum degree of all nodes at time-point *s*) denote the Newman-Girvan null model of intra-network connectivity. As this multi-layer modularity algorithm only allows positive matrix values, we removed all negative correlations in the sliding-window matrices, *A*. *γ*_*s*_ is the topological resolution parameter of time-point, or ‘layer’, *s*, and *ω* _*jrs*_ is the temporal coupling parameter for node *j* between time-point *r* and *s*. Then, *δ* (*M*_*is*_ *M*_*js*_) and *δ* (*M*_*is*_ *M*_*jr*_) are 1 if nodes belong to in the same module, and 0 if they do not belong to the same module (*M*). This process was, on average, iterated 5 times before the inherent heuristics of the multilayer modularity algorithm converged. Networks had an average *Q*-value of 0.59 ± 0.012 s.d., and an average of 3 modules per subject (range: 2-6 modules). The final output of the multilayer modularity algorithm was a two-dimensional array (*N*×*W*) with integer values denoting modules with strong within-network connectivity. The switching rate for each node was then estimated as the percentage of time-windows when a brain node transitions between different network assignments.

As discussed previously, it is non-trivial to select *γ* and *ω* parameters and we used a range of parameters including *γ* = [0.9, 1, 1.1] and *ω* = [0.5, 0.75, 1]. As shown in Supplementary Figure 5, the temporal *γ* parameter appeared to alter spatiotemporal modularity more than the topological *ω* parameter. Specifically, lower *ω* values led to increased network switching. Nodes switched 1.48% of time for *ω* = 1; 1.55% of time for *ω* = 0.75; and 1.61% of the time for *ω* = 0.5, at a constant *γ* value of 1.

### fMRI dynamic connectivity

The standard deviation of fMRI sliding-windowed correlation time-series between node pairs was here used as a proxy of dynamic connectivity, where high standard deviation indicates greater signal dispersion from mean correlation-based sliding window time-series. To determine whether our obtained standard deviation values of time-resolved fMRI connectivity likely reflect ‘true dynamics’ (meaning that we obtain information from this measure that cannot be obtained in time-averaged, or static, analysis), we compared standard deviation between original and phase randomised data where fMRI time-series were phase shuffled in the Fourier domain while preserving the power spectral magnitude and the correlational nature of the data (Prichard & Theiler, 1994). We obtained 500 phase randomized signals and used False Discovery Rate (*q* = 0.05) to reduce probability of type-I errors given that each subject has 300 unique comparisons. In total, 28.9 % of node-pairs were deemed ‘dynamic’ after correcting for multiple comparisons. To convert the dynamic connectivity data from matrix ([(*N*(*N* - 1)*/*2)] = 300) to node (*N* = 25) space, we summarized the (binary) number significant connections for each of, resulting in a degree metric summarized how ‘dynamic’ a node is.

### Sample Entropy

We used Sample Entropy to estimate the difference in temporal complexity of time-resolved fMRI connectivity signals between 5 nodes with most network switching and 5 nodes with least network switching. We used a epoch length, *M*, of 2 and a tolerance parameter, *r*, of 0.2×the standard deviation of the signal. See (Richman & Moorman, 2000), for more information about Sample Entropy. In brief, a low Sample Entropy score suggests a signal is regular or whereas a high Sample Entropy score suggests the signal is random or unpredictable.

### Cross-validated Elastic net regression

We used Elastic net regression to test whether whole-brain averaged network switching predicted 50 behavioral variables across subjects. Elastic net enables data-driven regression analysis by enforcing sparsity of regression output values (i.e., reducing the number of final *β* regression values). In other words, it provides automatic variable selection by removing all behavioral variables not predicted by network switching.

Given that network switching data and behavioral data had different numerical scales, we normalized all input data, *x*, which denotes both switching data and the 50 behavioral variables.

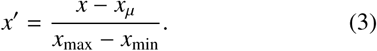

This resulted in variables, *x*′, with values between −1 and 1. The Elastic net equation is then written as (*y* = a vector of size 1×1003 containing subject-specific information from whole brain averaged network switching data; *X* = an array of size 50×1003 containing subject-specific information from 50 behavioral variables):

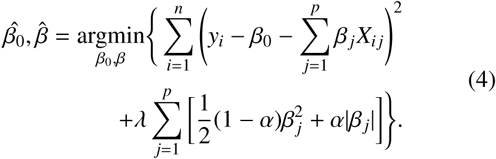

This is a doubly penalized regression model using both LASSO regression (*α* = 0; an *l*-1 penalty resulting in a sparse but uncorrelated *β* values) (Tibshirani, 1996) and Ridge regression (*α* = 1; an *l*-2 penalty resulting in a variance-reducing, but non-sparse *β* values) (Hoerl & Kennard, 1970). We set the *α* value to 0.5 to take advantage of the relative strengths of the two above regression approaches, providing a non-sparse solution with low-variance among several correlated behavioural independent variables (Supplementary Figure 6).

To select a *λ* threshold, which determines the overall sparsity of the regression model, we calculated Elastic net’s over a range of different *λ* values between 0 and 1 with increments of 0.001 (total of 1001 *λ* values) using 10-fold cross-validation (≈ 900 people were trained and ≈ 100 people were left out for testing, repeated 10 times until all subject have been left out once for training). The ‘optimal’ threshold had lowest mean square error over all possible *λ* ‘s across the 10-folds. We found *λ* = 0.023 had the lowest mean square error of 0.028 (see Supplementary Figure 3).

As reported in the main text, our results showed network switching predicted 3 of 50 behavioral variables. We defined prediction as:

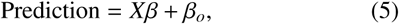

where X is the original matrix values of our 3 behavioural variables and *β*_*o*_ is the intercept of the Elastic net regression model.

## Acknowledgements

All fMRI data in this study has been made freely available by the Human Connectome Project, WUMinn Consortium (1U54MH091657; Principal Investigators: David Van Essen and Kamil Ugurbil) funded by the 16 National Institutes of Health (NIH) institutes and centers that support the NIH Blueprint for Neuroscience Research; and by the McDonnell Center for Systems Neuroscience at Washington University.

We acknowledge support from the National Health and Medical Research Council (NHMRC) of Australia (APP628952). The Florey Institute of Neuroscience and Mental Health acknowledges the strong support from the Victorian Government and in particular the funding from the Operational Infrastructure Support Grant. We also acknowledge the facilities, and the scientific and technical assistance of the National Imaging Facility (NIF) at the Florey node and The Victorian Biomedical Imaging Capability (VBIC). G Jackson is supported by the NHMRC practitioner’s fellowship (APP1060312). A Zalesky is supported by the NHMRC Senior Research Fellowship B (APP1136649).

## Appendix

**Figure S1.**
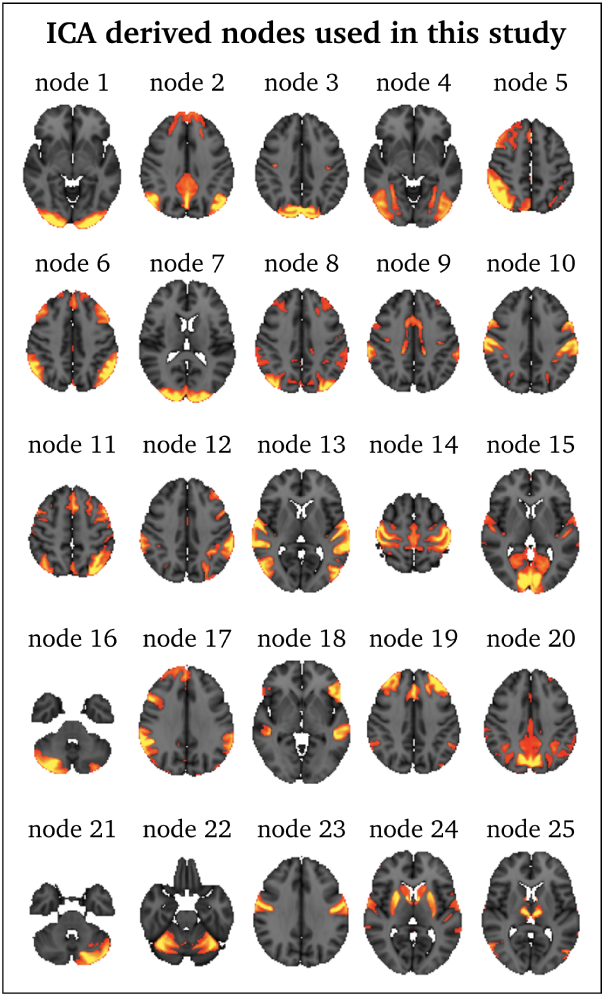
Brain nodes used in this study

**Figure S2.**
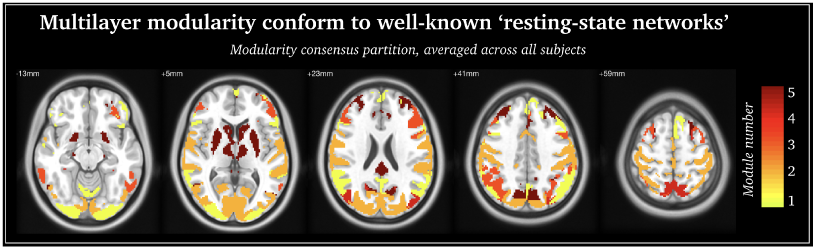
Group-consensus partition based on multilayer network modularity model – module number 1: frontoparietal network; module number 2: primary sensory cortices; module number 3: secondary sensory network; module number 4: fronto-parietal network; module number 5: salience/subcorical network.

**Figure S3.**
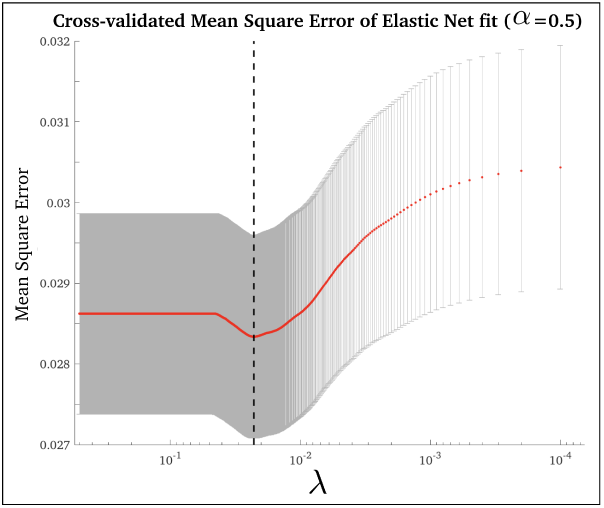
Mean square error -y-axis-of the 10-fold cross-validated Elastic net regression, across a range of λ -x-axis-

**Figure S4.**
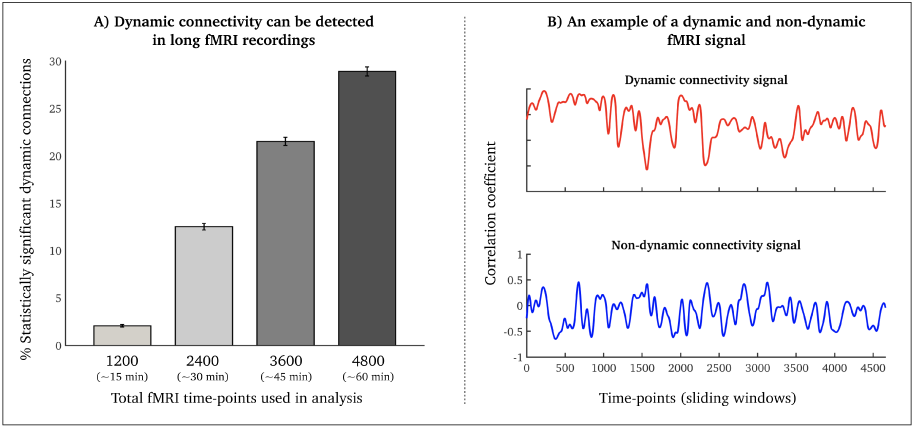
A) Increased probability of statistically significant connectivity dynamics in long fMRI recordings. B) examples of dynamic/non-dynamics fMRI connectivity signals

**Figure S5.**
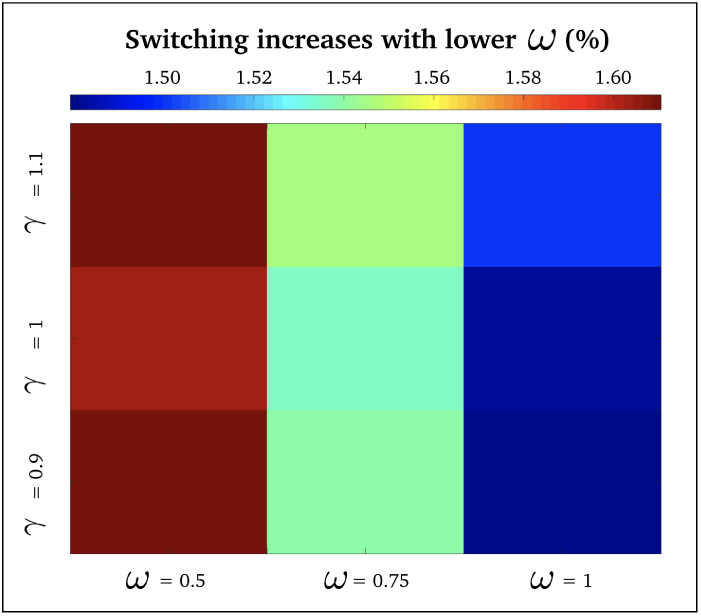
Effects of different spatial γ (y-axis) and temporal ω (x-axis) on network switching (color-scale)

**Figure S6.**
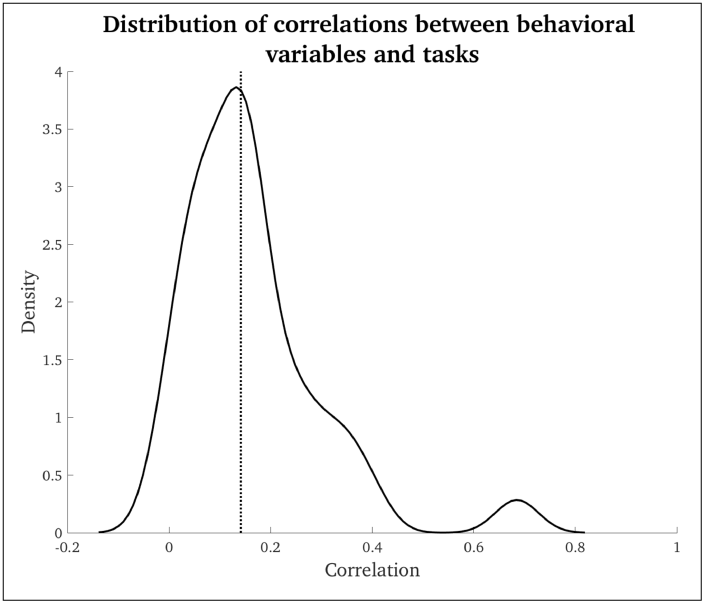
Distribution of correlations between task performance across subjects (50 tasks and conditions)

## References

Alhola, P., & Polo-Kantola, P. (2007). Sleep deprivation: Impact on cognitive performance. Neuropsychiatric disease and treatment.

Amigó, J. M., Kloeden, P. E., & Giménez, Á. (2013). Entropy increase in switching systems. Entropy, 15(6), 2363–2383.

Bandettini, P. A., Wong, E. C., Hinks, R. S., Tikofsky, R. S., & Hyde, J. S. (1992). Time course epi of human brain function during task activation. Magnetic resonance in medicine, 25(2), 390–397.

Barch, D. M., Burgess, G. C., Harms, M. P., Petersen, S. E., Schlaggar, B. L., Corbetta, M., … others (2013). Function in the human connectome: task-fmri and individual differences in behavior. Neuroimage, 80, 169–189.

Bassett, D. S., Porter, M. A., Wymbs, N. F., Grafton, S. T., Carlson, J. M., & Mucha, P. J. (2013). Robust detection of dynamic community structure in networks. Chaos: An Interdisciplinary Journal of Nonlinear Science, 23(1), 013142.

Bassett, D. S., Wymbs, N. F., Porter, M. A., Mucha, P. J., Carlson, J. M., & Grafton, S. T. (2011). Dynamic reconfiguration of human brain networks during learning. PNAS, 108(18), 7641–7646.

Benjamini, Y., & Hochberg, Y. (1995). Controlling the false discovery rate: a practical and powerful approach to multiple testing. Journal of the royal statistical society. Series B (Methodological), 289–300.

Betzel, R. F., Satterthwaite, T. D., Gold, J. I., & Bassett, D. S. (2017). Positive affect, surprise, and fatigue are correlates of network flexibility. Scientific Reports, 7(1), 520.

Braun, U., Schäfer, A., Walter, H., Erk, S., Romanczuk-Seiferth, N., Haddad, L., … others (2015). Dynamic reconfiguration of frontal brain networks during executive cognition in humans. PNAS, 112(37), 11678–11683.

Bullmore, E., & Sporns, O. (2009). Complex brain networks: graph theoretical analysis of structural and functional systems. Nature Reviews Neuroscience, 10(3), 186.

Courtney, S. M., Petit, L., Maisog, J. M., Ungerleider, L. G., & Haxby, J. V. (1998). An area specialized for spatial working memory in human frontal cortex. Science, 279(5355), 1347–1351.

De Domenico, M. (2017). Multilayer modeling and analysis of human brain networks. GigaScience, 6(5), 1–8.

Glasser, M. F., Sotiropoulos, S. N., Wilson, J. A., Coalson, T. S., Fischl, B., Andersson, J. L., … others (2013). The minimal preprocessing pipelines for the human connectome project. Neuroimage, 80, 105–124.

Hindriks, R., Adhikari, M. H., Murayama, Y., Ganzetti, M., Mantini, D., Logothetis, N. K., & Deco, G. (2016). Can sliding-window correlations reveal dynamic functional connectivity in resting-state fmri? Neuroimage, 127, 242–256.

Hoerl, A. E., & Kennard, R. W. (1970). Ridge regression: Biased estimation for nonorthogonal problems. Technometrics, 12(1), 55–67.

Hutchison, R. M., Womelsdorf, T., Allen, E. A., Bandettini, P. A., Calhoun, V. D., Corbetta, M., … others (2013). Dynamic functional connectivity: promise, issues, and interpretations. Neuroimage, 80, 360–378.

Keilholz, S. D., Magnuson, M. E., Pan, W.-J., Willis, M., & Thompson, G. J. (2013). Dynamic properties of functional connectivity in the rodent. Brain connectivity, 3(1), 31–40.

Kwong, K. K., Belliveau, J. W., Chesler, D. A., Goldberg, I. E., Weisskoff, R. M., Poncelet, B. P., … Turner, R. (1992). Dynamic magnetic resonance imaging of human brain activity during primary sensory stimulation. PNAS, 89(12), 5675–5679.

Lachenbruch, P. A., & Mickey, M. R. (1968). Estimation of error rates in discriminant analysis. Technometrics, 10(1), 1–11.

Lancichinetti, A., & Fortunato, S. (2012). Consensus clustering in complex networks. Scientific reports, 2, 336.

Leonardi, N., & Van De Ville, D. (2015). On spurious and real fluctuations of dynamic functional connectivity during rest. Neuroimage, 104, 430–436.

Mucha, P. J., Richardson, T., Macon, K., Porter, M. A., & Onnela, J.-P. (2010). Community structure in time-dependent, multiscale, and multiplex networks. science, 328(5980), 876–878.

Muldoon, S. F., & Bassett, D. S. (2016). Network and multilayer network approaches to understanding human brain dynamics. Philosophy of Science, 83(5), 710–720.

Ogawa, S., Tank, D. W., Menon, R., Ellermann, J. M., Kim, S. G., Merkle, H., & Ugurbil, K. (1992). Intrinsic signal changes accompanying sensory stimulation: functional brain mapping with magnetic resonance imaging. PNAS, 89(13), 5951–5955.

Pedersen, M., Omidvarnia, A., Zalesky, A., & Jackson, G. D. (2018). On the relationship between instantaneous phase synchrony and correlation-based sliding windows for time-resolved fmri connectivity analysis. NeuroImage, 181, 85–94.

Preti, M. G., Bolton, T. A., & Van De Ville, D. (2017). The dynamic functional connectome: State-of-the-art and perspectives. Neuroimage, 160, 41–54.

Prichard, D., & Theiler, J. (1994). Generating surrogate data for time series with several simultaneously measured variables. Physical review letters, 73(7), 951.

Richman, J. S., & Moorman, J. R. (2000). Physiological time-series analysis using approximate entropy and sample entropy. American Journal of Physiology-Heart and Circulatory Physiology, 278(6), H2039–H2049.

Rubinov, M., & Sporns, O. (2010). Complex network measures of brain connectivity: uses and interpretations. Neuroimage, 52(3), 1059–1069.

Salimi-Khorshidi, G., Douaud, G., Beckmann, C. F., Glasser, M. F., Griffanti, L., & Smith, S. M. (2014). Automatic denoising of functional mri data: combining independent component analysis and hierarchical fusion of classifiers. Neuroimage, 90, 449–468.

Shine, J. M., Koyejo, O., & Poldrack, R. A. (2016). Temporal metastates are associated with differential patterns of time-resolved connectivity, network topology, and attention. PNAS, 113(35), 9888–9891.

Telesford, Q. K., Ashourvan, A., Wymbs, N. F., Grafton, S. T., Vettel, J. M., & Bassett, D. S. (2017). Cohesive network reconfiguration accompanies extended training. Human brain mapping, 38(9), 4744–4759.

Tibshirani, R. (1996). Regression shrinkage and selection via the lasso. Journal of the Royal Statistical Society. Series B (Methodological), 267–288.

Van Den Heuvel, M. P., & Sporns, O. (2011). Rich-club organization of the human connectome. Journal of Neuroscience, 31(44), 15775–15786.

Van Essen, D. C., Smith, S. M., Barch, D. M., Behrens, T. E., Yacoub, E., Ugurbil, K., … others (2013). The wu-minn human connectome project: an overview. Neuroimage, 80, 62–79.

Van Essen, D. C., Ugurbil, K., Auerbach, E., Barch, D., Behrens, T., Bucholz, R., … others (2012). The human connectome project: a data acquisition perspective. Neuroimage, 62(4), 2222–2231.

Wager, T. D., & Smith, E. E. (2003). Neuroimaging studies of working memory. Cognitive, Affective, & Behavioral Neuroscience, 3(4), 255–274.

Zalesky, A., & Breakspear, M. (2015). Towards a statistical test for functional connectivity dynamics. Neuroimage, 114, 466–470.

Zalesky, A., Fornito, A., Cocchi, L., Gollo, L. L., & Breakspear, M. (2014). Time-resolved resting-state brain networks. PNAS, 111(28), 10341–10346.

Zheng, H., Li, F., Bo, Q., Li, X., Yao, L., Yao, Z., … Wu, X. (2017). The dynamic characteristics of the anterior cingulate cortex in resting-state fmri of patients with depression. Journal of affective disorders, 227, 391–397.

Zou, H., & Hastie, T. (2005). Regularization and variable selection via the elastic net. Journal of the Royal Statistical Society: Series B (Statistical Methodology), 67(2), 301–320.

